# Potent neutralization of 2019 novel coronavirus by recombinant ACE2-Ig

**DOI:** 10.1101/2020.02.01.929976

**Authors:** Changhai Lei, Wenyan Fu, Kewen Qian, Tian Li, Sheng Zhang, Min Ding, Shi Hu

## Abstract

2019-nCoV, which is a novel coronavirus emerged in Wuhan, China, at the end of 2019, has caused at least infected 11,844 as of Feb 1, 2020. However, there is no specific antiviral treatment or vaccine currently. Very recently report had suggested that novel CoV would use the same cell entry receptor, ACE2, as the SARS-CoV. In this report, we generated a novel recombinant protein by connecting the extracellular domain of human ACE2 to the Fc region of the human immunoglobulin IgG1. An ACE2 mutant with low catalytic activity was also used in the study. The fusion proteins were then characterized. Both fusion proteins has high affinity binding to the receptor-binding domain (RBD) of SARS-CoV and 2019-nCoV and exerted desired pharmacological properties. Moreover, fusion proteins potently neutralized SARS-CoV and 2019-nCoV *in vitro.* As these fusion proteins exhibit cross-reactivity against coronaviruses, they could have potential applications for diagnosis, prophylaxis, and treatment of 2019-nCoV.

---

2019-nCoV, which is a novel coronavirus emerged in Wuhan, China, at the end of 2019, has caused at least infected 11,844 as of Feb 1, 2020. However, there is no specific antiviral treatment or vaccine currently. Spike (S) proteins of coronaviruses, including the coronavirus that causes severe acute respiratory syndrome (SARS) and the 2019-nCoV, associate with cellular receptors to mediate infection of their target cells. Angiotensin-converting enzyme 2 (ACE2), a carboxypeptidase that potently degrades angiotensin II to angiotensin 1–7, has been identified as a functional receptor for SARS-CoV^1^ and a potent receptor for 2019-nCoV^2^. This metallopeptidase is also playing key role in the renin-angiotensin system (RAS)^3^.

As ACE2 is a key player in the hormonal cascades that plays a central role in the homeostatic control of cardiorenal actions, earlier researchers had exerted effect on the pharmacodynamics studies of recombinant ACE2 (rACE2) to protect against severe acute lung injury and acute Ang II-induced hypertension^4-6^. In mice models, administration of rACE2 also showed to adverse myocardial remodelling, attenuate Ang II-induced heart hypertrophy and cardiac dysfunction,^7^, as well as renal oxidative stress, inflammation, and fibrosis.^8,9^ However, both in humans and mice^10,11^, rACE2 exhibits fast clearance rates with a reported half-life of only hours by pharmacokinetic studies. Recently, fusion protein of rACE2 with an Fc fragment (rACE2-Fc) showed organ protection in both acute and chronic models of angiotensin II-dependent hypertension in mice^12^, interestingly, the fusion protein also showed long-lasting effects.

Based on the receptor function of ACE2 for coronavirus, we hypothesis that ACE2 fusion protein may have neutralization potential for coronavirus, especially the 2019-nCoV. To further evaluated the therapeutic potential of ACE2, fusion protein (ACE2-Ig) consisting of the extracellular domain of human ACE2 linked to the Fc domain of human IgG1 was constructed and generated (Fig. 1). An ACE2 variant (HH/NN) in which two active-site histidines (residues 374 and 378) have been altered to asparagines was also used in our study to reduce the catalytic activity. The fusion protein contain the ACE2 variant were termed as mACE2-Ig. The expression and purification methods were described in our previous reports^13,14^. The affinities of the fusion proteins for SARS-CoV RBD and 2019-nCoV RBD were determined with BIAcore binding assays (Fig. S1) and both fusion proteins showed high affinity to RBDs. Moreover, the fusion proteins had similar denaturation temperature compared with the Fc fusion protein TIGIT-Ig reported in our pervious report^13^, suggesting a IgG-like stability. The lowest concentration (< 2%) of high molecular weight and low molecular weight products was observed after 1 week of storage at 40 °C at a 1 mg/ml concentration. Pharmacokinetic (PK) parameters of the fusion proteins were then tested in our study. Mice were treated separately with a single intravenous dose of fusion proteins and the serum concentrations of fusion proteins were determined by ELISA. The results showed that the main PK parameters of ACE2-Ig, mACE2-Ig and TIGIT-Ig were very similar in mice and demonstrated the high stability of the fusion proteins. The experimental data are summarized in Table S1.

**Figure 1.**
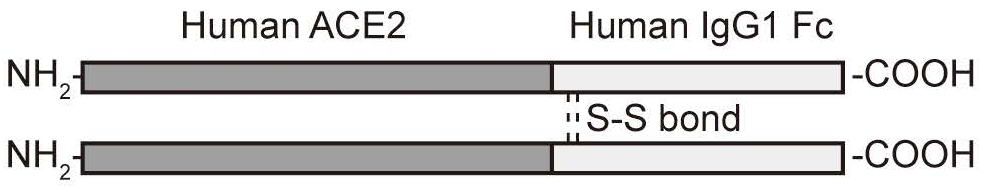
Schematic of ACE2-Ig.

After we identified that ACE2 fusion proteins binds to the RBDs with high affinity, we next sought to test the inhibitory activity of ACE2 fusion proteins against 2019-nCoV and compare it with that against SARS-CoV, by using viruses pseudotyped with the S glycoprotein of SARS-CoV and 2019-nCoV. Our data shows that Both SARS-CoV and 2019-nCoV viruses were potently neutralized by ACE2-Ig and mACE2-Ig. The IC50 of SARS-CoV and 2019-nCoV viruses neutralized by ACE2-Ig were 0.8 and 0.1 μg/ml, respectively. And The IC50 of the two viruses neutralized by mACE2-Ig were 0.9 and 0.08 μg/ml, respectively (Fig. 2). No evidence of neutralization was observed for the TIGIT-Ig. We next using cell fusion assay to further characterize the in vitro neutralization effect of the fusion proteins (Fig. 3). ACE2-Ig potently inhibited the SARS CoV-S protein-mediated fusion with an IC50 of 0.85 μg/ml, and the 2019 nCoV-S protein-mediated fusion with an IC50 of 0.65 μg/ml. Under the same experimental conditions, another fusion protein mACE2-Ig exhibited an IC50 of 0.76 μg/ml and 0.48μg/ml, for the SARS CoV and 2019 nCoV, respectively. The control TIGIT-Fc did not show any inhibitory effect in this assay. These data suggest that both ACE2-Ig and mACE2-Ig exhibit potent inhibitory activity against SARS CoV and 2019 nCOV.

**Figure 2.**
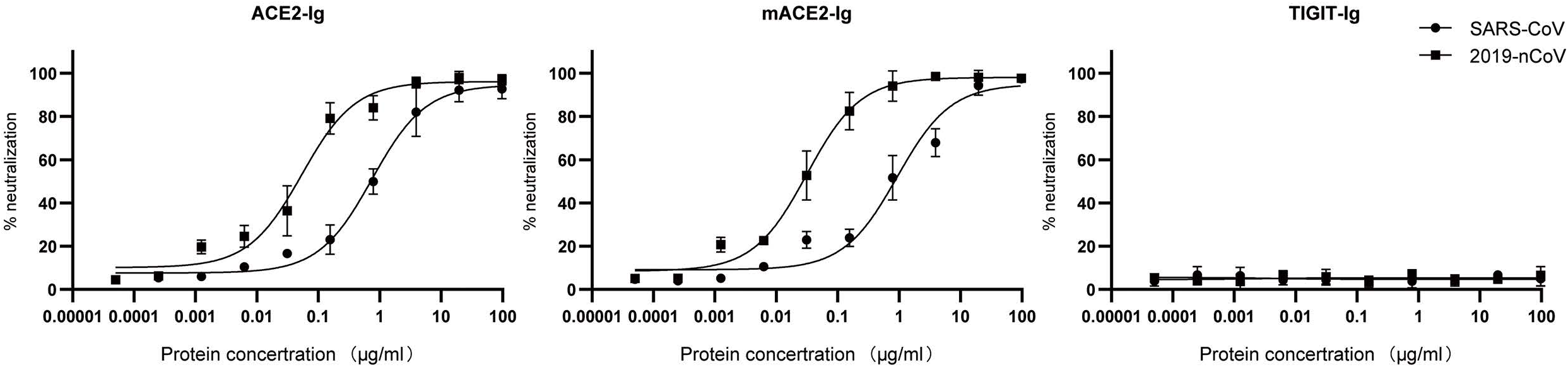
ACE2-Ig potently neutralizes viruses pseudotyped with S glycoproteins from the SARS CoV and 2019 nCoV. HIVs pseudotyped with the S glycoprotein from CoVs were incubated with different fusion proteins for 1 h before infection. Luciferase activities in target cells were measured, and the percent neutralization was calculated.

**Figure 3.**
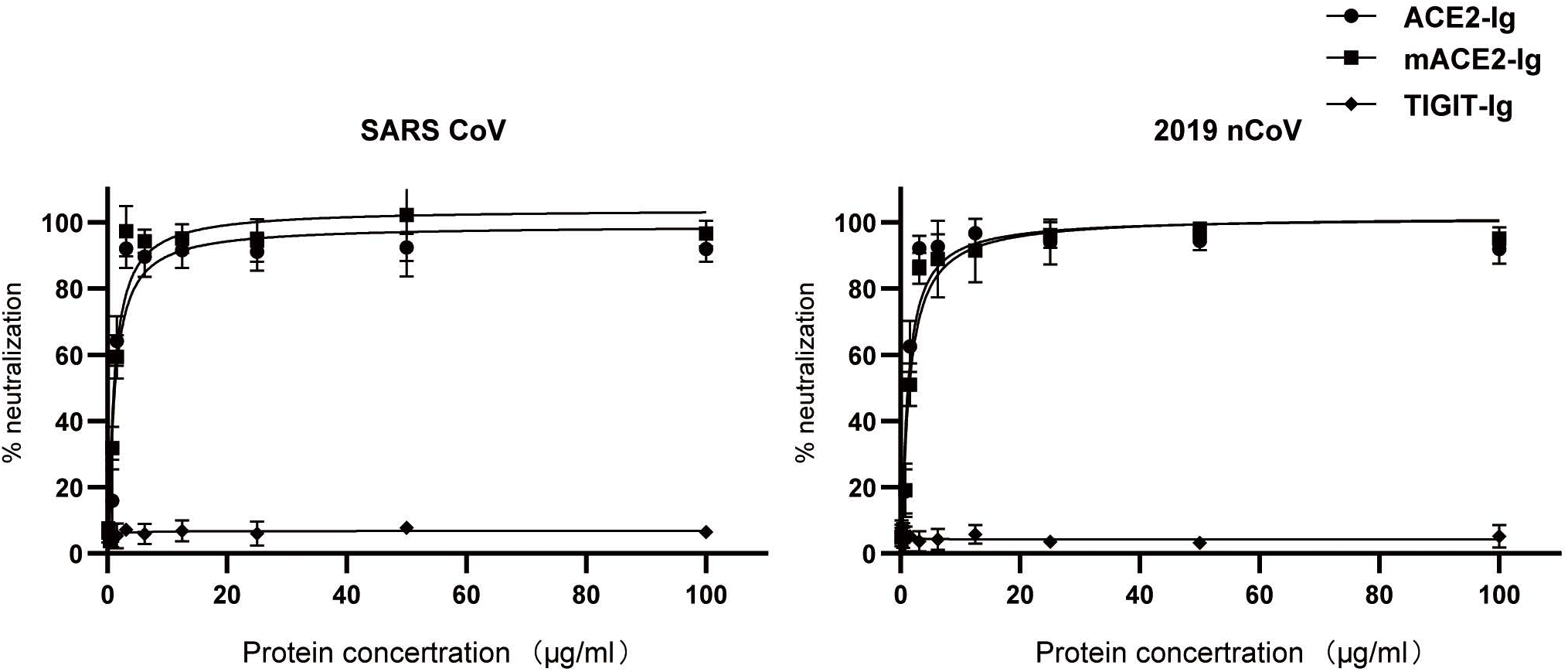
Inhibition of cell fusion by ACE2-Ig. Potent inhibition of cell fusion mediated by the SARS-CoV (left) or 2019 nCoV (right) spike (S) glycoprotein. Cells expressing diffferent S glycoprotein were incubated with indicated fusion proteins and mixed with ACE2-expressing cells. The activity of the reporter gene, β-gal, was measured as a correlate of fusion. The curves represent the best fit to the experimental data and were used for calculation of the IC50. Bars indicate the S.E.

ACE2 has already been identified as an important drug targets for the treatment of cardiovascular and kidney diseases. Although it has been established that ACE2 is a key function receptor for the coronavirus infection, the practical use of rACE2 protein may be impede by the short half-life of rACE2. Recombinant Fc fusion technique, which enable soluble proteins to extend plasma residence time, improve in vivo efficacy, and gain immunoreactive functions, has been widely used in modern biopharmaceuticals. For example, new long-acting forms of both recombinant coagulation factors rFVIII-Fc and rFIX-Fc were recently approved for clinical treatments of hemophilia A and B that require less frequent infusions^15,16^. And the TNF receptor fusion protein has been use in clinic for decades^17^. Unlike the blood resident enzymes of coagulation factors, full-length endogenous ACE2 is a transmembrane protein anchored to the cell surface and that ACE2 activities are, in fact, present at very low levels in systemic circulation^18-20^. Therefore, one safety concern of the ACE2 fusion protein is that they may have systemic cardiovascular side-effect. Interestingly, one recently report shows that treatment of murine ACE2 fusion proteins in mice show no evidence of side-effect^12^. Moreover, our preliminary studies showed that the neutralization effect remained efficient when two active-site histidines of ACE2 were modified to asparagine.

To sum up, our data investigate the therapeutic potential of ACE2 based strategies against 2019-nCoV which could be used alone or in combination. Based on the molecular mechanisms of blocking of ACE2, the fusion proteins have the potential to have broad neutralizing activity to coronavirus. These ACE2 fusion proteins could be also used for diagnosis and as research reagents in the development of vaccines and inhibitors.

## Methods

### Generation of fusion proteins

For recombinant fusion plasmid, DNA sequences of extracellular domains (ECDs) of ACE2 (aa: 1-740) were ligated to the Fc segment of human IgG1. Mutations were generated by integrated DNA Technologies. We use the FreeStyle 293 expression system (Invitrogen) to express the fusion proteins. Vectors and detailed methods were according to our previous study^21^. Fusion proteins were further purified using protein A Sepharose with the harvested cell culture supernatant. TIGIT-Ig, which were described in our pervious report^13^, served as a control in our study. The concentration and purity of the fusion protein were determined by measuring the UV absorbance at a wavelength of 280 nm and by polyacrylamide gel electrophoresis, respectively.

### Affinity measurement

CoV RBDs were prepared as previously described^1^. We use a previously reported method^14^ with immobilized anti-human Fc polyclonal antibody on a CM5 chip (∼150 RU) to capture the fusion proteins and and then injected CoV RBDs (12.5 nM∼200 nM). blank flow cells were used for the correction of the binding response. The surface plasmon resonance (SPR) method with a BIAcore-2000 was used to measure the monovalent binding affinity of the fusion protein and performed kinetic analysis using a 1:1 L model that simultaneously fit ka and kb.

### Pharmacokinetics

We used BALB/c mice to determine the pharmacokinetic profile of the fusion protein. Eight-week-old mice were administered the fusion protein at a dose of 5 mg/kg body weight by tail vein injection. Mice were divided into 15 groups, corresponding to day 1 to day 15. Blood was collected from the septum in heparin-containing tubes and then centrifuged to remove blood cells and to obtain plasma samples. The serum concentration of the fusion protein was determined by ELISA.

### Pseudovirus Neutralization Assay

Pseudoviruses containing the S glycoprotein from various virus, and a defective HIV-1 genome that expresses luciferase as a reporter protein, were prepared, and the assays performed as described previously^22-25^.

### Cell Fusion Inhibition Assay

A quantitative cell fusion assay based on β-galactosidase (β-gal) as a reporter gene was used for the assessment of the neutralization activity of the fusion proteins, as described previously^26^. 293T cells transfected with the indicated CoV S glycoprotein genes were preincubated with different fusion proteins at room temperature for 15 min, then mixed with 293T cells transfected with ACE2 at 1:1 ratio and incubated at 37°C for 4 h. Cells were then lysed, and the β-gal activity was measured. The protein concentrations during fusion were used for calculation of the IC50 defined as the concentration at which the β-gal activity was reduced by 50%.

