## Supplementary Figures and Table for "Potent neutralization of 2019 novel coronavirus by recombinant ACE2-Ig"

**Contents** **Page**

### Supplementary Figures

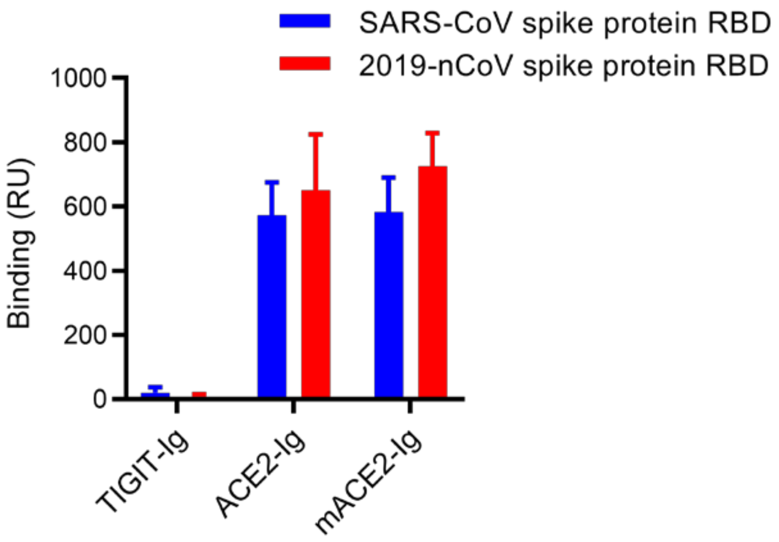

**Figure S1. Binding of Fusion proteins to CoV spike protein RBDs**

Fusion proteins was tested for binding to immobilized CoV spike protein RBDs using surface plasmon resonance on a BIAcore 2000 instrument. Binding was quantified as an increase in RU at 60 s after the end of injection compared with a baseline established 20 s before injection.

### Supplementary Tables

#### Table S1. Selected analytical data and pharmacokinetic parameters of recombinant fusion proteins in mice.

| Parameter[a] | ACE-Ig | mACE2-Ig | TIGIT-Ig |
|---|---|---|---|
| HMW formation after storage(% SEC area)[b] | < 0.1 | < 0.1 | < 0.1 |
| LMW formation after storage(% SEC area)[b] | < 0.1 | < 0.1 | < 0.1 |
| AUC (day µg $ml^{-1}$) | 540.8 | 464.3 | 557.2 |
| $T_{1/2}$(day) | 5.2 | 5.3 | 4.7 |
| CL (ml $day^{-1}$ $kg^{-1}$) | 7.2 | 8.5 | 7.3 |
| VSS(ml $kg^{-1}$) | 68.3 | 78.0 | 65.6 |

[a] Pharmacokinetic parameters were calculated using a noncompartmental analysis. AUC, area under the concentration versus time curve; t1/2, half-life; CL, clearance; VSS, steady-state volume of distribution.
[b] Quiescent storage for 1 wk, 40 °C, 1 mg/mL
